# Decentralized Brain Age Estimation using MRI Data

**DOI:** 10.1101/2021.05.10.443469

**Authors:** Sunitha Basodi, Rajikha Raja, Bhaskar Ray, Harshvardhan Gazula, Jingyu Liu, Eric Verner, Vince D. Calhoun

## Abstract

Recent studies have demonstrated that neuroimaging data can be used to predict brain age, as it captures information about the neuroanatomical and functional changes the brain undergoes during development and the aging process. However, researchers often have limited access to neuroimaging data because of its challenging and expensive acquisition process, thereby limiting the effectiveness of the predictive model. Decentralized models provide a way to build more accurate and generalizable prediction models, bypassing the traditional data-sharing methodology. In this work, we propose a decentralized method for brain age estimation and evaluate it on three different feature sets, including both volumetric and voxelwise structural MRI data as well as resting functional MRI data. The results demonstrate that a decentralized brain age model can achieve similar performance compared to the models trained with all the data in one location.

## 1 Introduction

Brain age estimation (BAE) from magnetic resonance imaging (MRI) data has become widely popular in recent years. Computed as the difference between chronological age and estimated brain age, the brain age gap is helpful in early identification of various neurodegenerative diseases [1, 2, 3] and was also found to be wider in patients with dementia and autism [4]. Several studies have used traditional machine learning [5, 6] and deep learning approaches [7] to develop promising prediction models on large datasets that have either been collected at one location or sourced from different locations. Although having a large training dataset can help in achieving robust models, there are limitations on the amount of data that can be gathered for such large-scale analyses. Examples of such limitations include the cost-prohibitiveness of MRI data collection, varying institutional data-sharing policies, constrained data-usage agreements, data-privacy concerns, and many more [8].

One efficient approach to bypass these limitations is to use decentralized algorithms, which do not require pooling the data in one location. Decentralized learning algorithms have become popular in recent years due to the demand for collaborative studies [8, 9]. Such approaches are particularly important when there is a need to perform a large-N analysis involving heterogeneous datasets without worrying about data transmission or violating privacy.

In this work, we have developed a decentralized brain age prediction algorithm using support vector regression and performed detailed experiments on three different feature sets (types of datasets) using varying sampling methods and demonstrate the robustness of our approach. We implement our approach within the Collaborative Informatics and Neuroimaging Suite Toolkit for Anonymous Computation (COINSTAC) [9] framework (more on this later). Results show that the proposed decentralized method attained performance on par with a centralized approach.

The remainder of this paper is organized as follows. The background on brain age, current prediction approaches, decentralized learning, and COINSTAC are presented in section 2. The decentralized BAE method is presented in section 3, followed by a discussion of the results in section 4 and conclusion in section 5.

## 2 Background

### 2.1 Brain age

The brain age (of an individual) is the observable age of the brain in contrast to the chronological (actual) age. As brain age cannot be directly measured, predictive models are typically trained with the chronological age of cognitively normal subjects. Any discrepancy between the estimated brain age and chronological age is considered to be the brain age gap. Studies have shown that patients suffering from neurodegenerative diseases (such as Alzheimer’s, Huntington’s, Parkinson’s, and Multiple Sclerosis) have an increased brain age gap or brain-predicted age [4]. In contrast, there are studies that show activities such as meditation [10] and physical exercise decrease brain age [11]. These and other studies [12, 13, 2] suggest BAE may serve as an important biomarker in understanding the aging process of the brain and its relationship to brain health and disorder.

### 2.2 Current prediction approaches

Predicting brain age on the basis of MRI is now a widely used approach. There are broadly three different approaches that have been previously used, namely, voxel-based, region-based, and surface-based approaches for structural MRI[14]. In a voxel-based approach, MR images are first segmented into gray matter (GM), white matter (WM), and cerebrospinal fluid (CSF), and then this voxel-level information is used to fit the models. In a region-based approach, an atlas is used to summarize the data for estimating prediction models and brain age. Surface-based methods generally compute a triangulated mesh using WM/GM boundaries or GM/CSF boundaries and then extract surface-based features for prediction. In this work, we also introduce a component-based approach for functional MRI (fMRI) data that uses the timecourses from independent components or rather their cross-correlations, called functional network connectivity (FNC), as features [15]. The different features extracted from each of these approaches can be used with either traditional machine learning or deep learning models. Deep learning models have shown promising results in BAE. In [12], BAE was computed using both raw MR images and preprocessed MR images, and both showed consistent results. The authors of [16] employ single and multimodal brain imaging data, including structural MRI (sMRI), diffusion tensor imaging (DTI), and resting state fMRI (rs-fMRI), and evaluate performance on many models. Recently, graph neural networks have also been used for BAE [17] using data from the UK Biobank [18]. However, there is no clear understanding about which models perform the best; though it is clear that having large amounts of training data usually helps in achieving robust models.

### 2.3 Decentralized learning

Decentralized learning has recently gained attention as it allows privacy-preserving, large-scale, machine learning analysis [19]. Also referred to as federated learning in literature, decentralized machine learning is focused on training models on data that are not shared but that researchers want to be analyzed together. In this method, all the participating sites start with the same model, i.e., a local model is trained on its own data. In each iteration, gradients from local models are sent back to the main site for aggregation, after which updated gradients are returned to local sites to update their models. This is repeated over many iterations until the model achieves its stopping criteria, such as the desired performance. There are several challenges in designing decentralized algorithms, including heterogeneity across different sites, transmission, maintaining synchronization during training iterations, and preserving data privacy. Heterogeneity arises from the different devices (hardware and software) involved during the training process, and one cannot overlook this issue, as it can effect the overall performance of a decentralized model. Though there is no raw data exchange, decentralized learning involves transmission of model gradients/parameters that utilize significant bandwidth, especially when training deep learning models. The main site also needs to ensure that the aggregation is performed on the correct iteration data from all the sites when there is repeated communication between the local and main site. For a detailed summary of the machine learning algorithms, challenges, and architectures for decentralization, we refer the readers to [20, 21].

### 2.4 COINSTAC

COINSTAC [8, 9, 22] is a platform that enables decentralized analysis of (neuroimaging) data without the need to pool the data at one location. It was created to enable collaborative research by removing the barriers to traditional data-centric approaches. As there is no pooling of data, COINSTAC protects the privacy of individual datasets and solves the problem of researchers wanting to collaborate but being unable to because of data sharing restrictions [23]. Gazula et. al [24] used this application to perform a decentralized regression on sMRI data preprocessed using voxel-based morphometry to analyze the structural changes in the brain as linked to age, body mass index, and smoking. This study also emphasized the benefits of large-scale neuroimaging analysis. COINSTAC implements a wide and growing range of decentralized neuroimaging pipelines and supports such large-scale analysis of decentralized data with results on par with results from pooled data.

## 3 Methods

Though decentralized learning has been applied in some domains, this paper, to our knowledge, presents the first approach for decentralized brain age analysis. In this work, we perform brain age prediction using a decentralized approach. Let us assume there are *N* + 1 participating sites, each gathering data from a different set of participants.

In centralized models, data from all the local sites is combined at a central location, where it is used for training a model, and therefore a centralized model has better performance than any model trained only on a subset of that data. In decentralized studies, instead of transferring the original data, only the information needed to train a model is shared, which not only improves the prediction performance of the model but also keeps the data secure at the local sites. Information sent from local sites is collected at the main site to build an aggregated model. The challenge is to reduce the performance gap between decentralized and centralized models without sharing the original data. Such a decentralized training approach can be used to train any prediction model; however, the type of information transferred between local and main sites highly depends on the type of the prediction model used (as different machine learning algorithms have different parameters and approaches to reach the optimal solution).

Formally, let N+1 be the total participating sites, where *site* – 0 is considered as main site and the remaining sites are local sites. Let the matrix 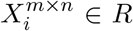 be data of *m* subjects with *n* features available at a site *i* ∈ {0,1,2,..,*N*}, with the matrix 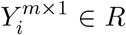 representing their chronological age. At each local site *i*, a regression model *M_i_* is constructed, and the corresponding model parameters 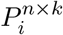 are sent to the main site, where *k* is the dimension of the parameters generated by the model.

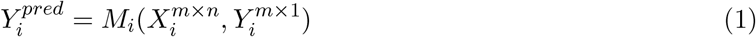

At main site (*site* – 0), these model parameters are aggregated and used to transform the original data 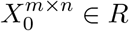. The data are transformed using the parameter matrix with the ⊗ operation. In our case, it is a simple matrix multiplication due to a linear regression model, although in general it can be any matrix operation. The transformed features 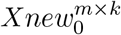 are used for BAE.

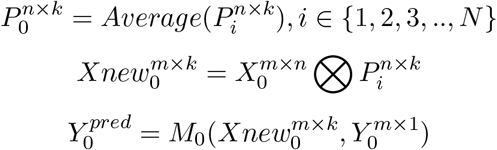

Here *M*_0_ represents the aggregated model containing information from models *M_i_*, *i* ∈ {1, 2,.., *N*}.

**Algorithm 1:**
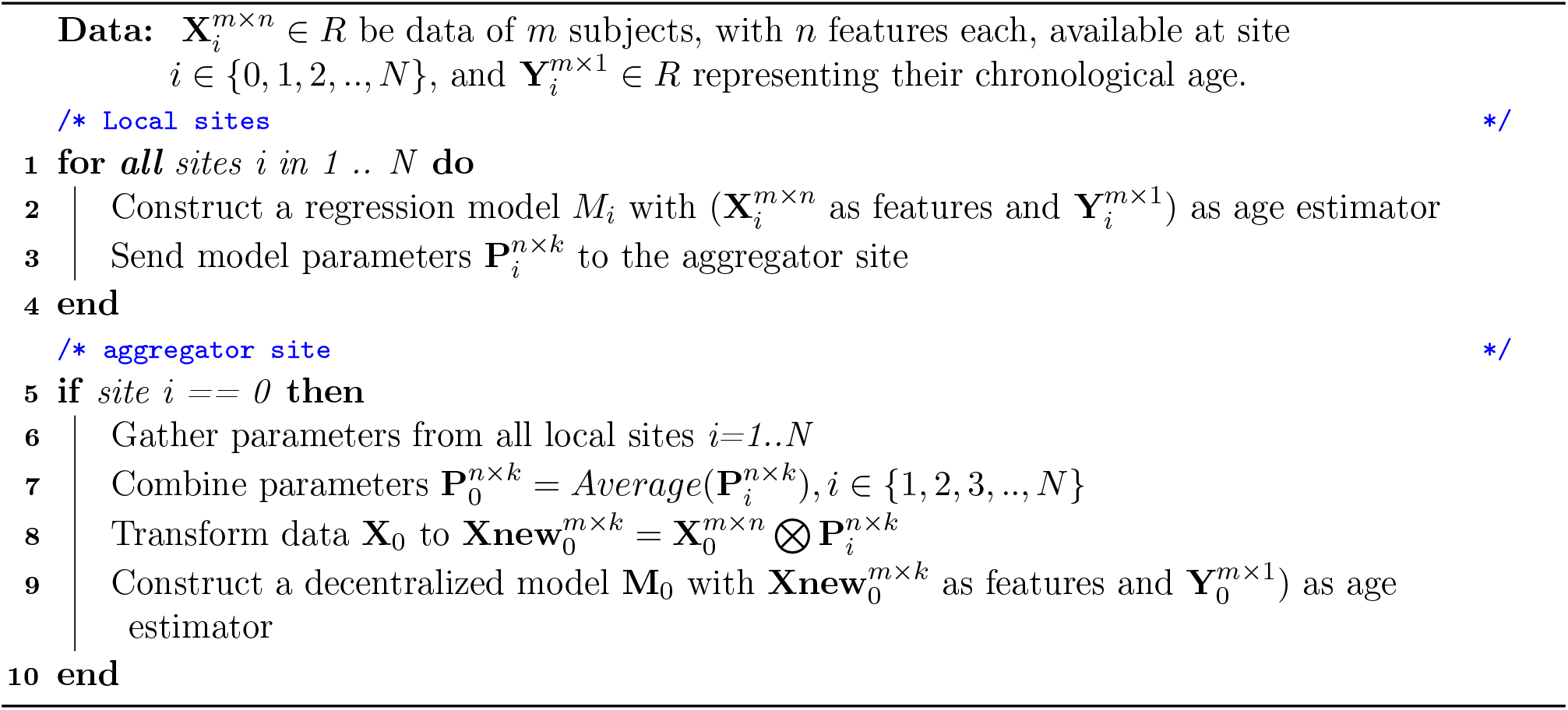
Decentralized brainage prediction

In our initial setup, all the local sites are provided with the details of the prediction model and the type of information to be shared with the main site after training their local models. Of all the participating sites, *N* sites are used as local sites, and the remaining site is used as the main site. In this study, we use support vector regression (SVR) as the prediction model and apply decentralization by employing a training strategy similar to [19, 25]. As a first step, all the local sites train an SVR model locally with their data and transfer the weight vectors of the locally learned models to the main site. The main site averages these vectors and uses the average weight vector to transform its data into a new feature space. This modified data is then used to train a decentralized SVR model (see Fig. 1).

**Figure 1:**
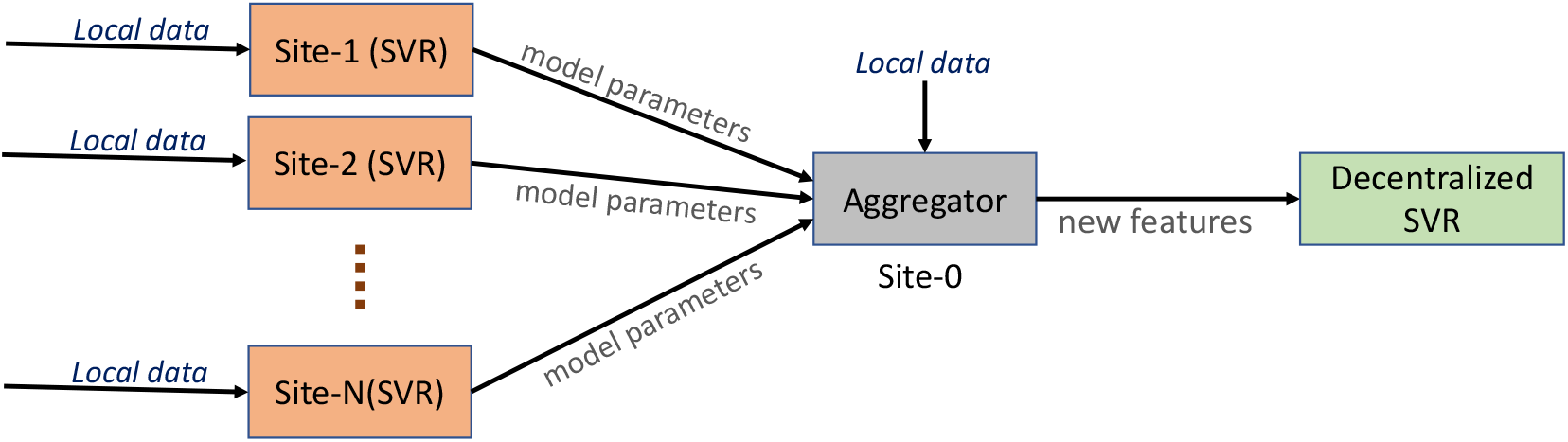
Overall flow of model parameters from locally trained SVR models that are sent to the main site for aggregation. The aggregated model parameters are used to transform the data at the main site into new features, which are used for training a new decentralized SVR model.

## 4 Results

The primary goal of this work is to develop a decentralized BAE model and contrast its performance against a centralized model. For this purpose, we performed three experiments with features extracted from two different datasets. All experiments were performed in COINSTAC [9] with six sites (one as main and the remaining five as local nodes).

### Datasets

There are two MRI datasets that are analyzed in this work, *viz*, UPENN-PNC [26] and UKBiobank [27]. The UPENN-PNC data consists of T1-weighted sMRI of 1417 health participants with chronological ages ranging from 8 to 21 years during acquisition. This data is jointly spatially normalized and segmented and then smoothed by a 6-mm full-width at half maximum (FWHM) Gaussian kernel. Segmentation was performed in SPM12 [28]. The second dataset, UKBiobank, has fMRI images of 11,754 subjects with chronological ages in the range of 44 to 80 years. The data were preprocessed using a combination of FSL [29] and SPM12. This included distortion correction, rigid body motion correction, normalization to Montreal Neurological Institute (MNI) space and smoothing using a Gaussian kernel with an FWHM of 6 mm. Next, we ran a fully-automated independent component analysis (ICA) pipeline that can capture corresponding functional network features while retaining more single-subject variability [30]. This pipeline has been successfully applied to multiple studies to identify a wide range of connectivity abnormalities in numerous brain diseases [30]. Group ICA was first performed on two large healthy control datasets to create spatial network (component) priors following which a spatially constrained ICA algorithm was applied to back-reconstruct spatial maps and time-courses (TCs) for each subject [31]. Fig. 2 shows the age range distribution of the subjects in these datasets. Fig. 3 shows the three different features extracted using these datasets as described below.

**Figure 2:**
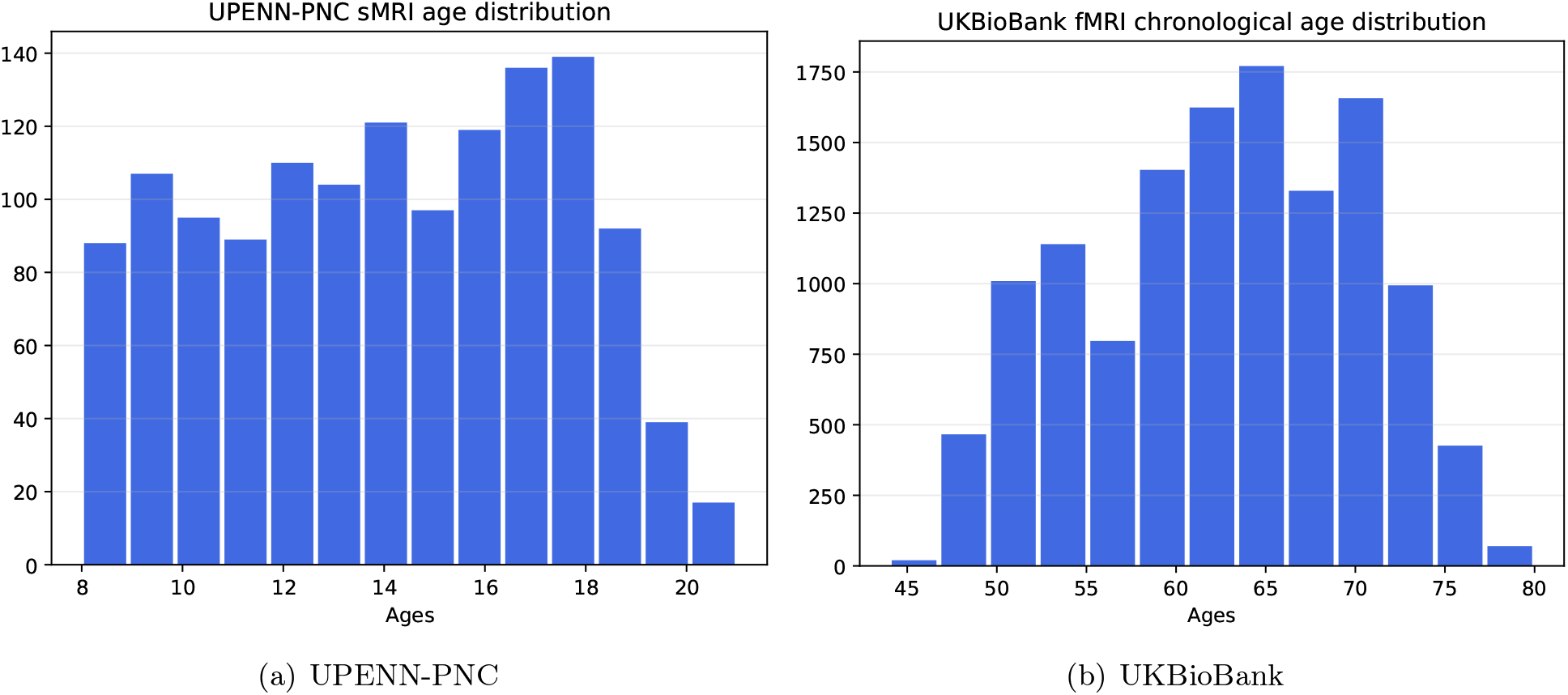
Histograms showing the chronological age distribution of healthy subjects of UPENN-PNC (left) and UKBiobank (right) datasets.

**Figure 3:**
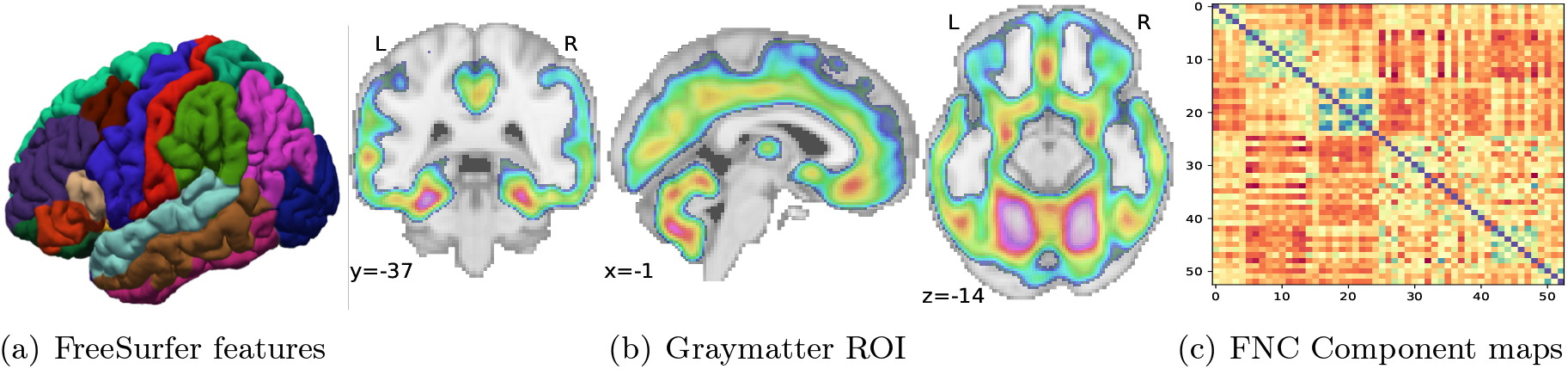
Different features extracted from sMRI and fMRI datasets for BAE.

### Prediction using FreeSurfer features

We used FreeSurfer (version v5.3) [32] to extract brain structural features from UPENN-PNC data. A standard `aseg.stats` file that has features corresponding to total Intracranial Volume (eTIV), left hemisphere (lh) and right hemisphere (rh) subcortical regions is generated. Additionally, features corresponding to cortical thicknesses and volumes of the parcellated regions in surface GM in both left and right hemispheres are also extracted. In total, we use 152 features for each subject to train the regression models.

### Prediction using GM features

We use the Automatic Anatomical Labeling (AAL) [33] brain atlas of 116 brain parcellations to extract the mean GM density of each region of interest (ROIs)from UPENN-PNC data for 1417 subjects. The extracted mean GM density of 116 ROIs are used as GM features to train the regression models.

### Prediction using FNC features

From the UKBiobank, we applied a fully automated spatially constrained ICA approach to fMRI data from 11,754 subjects, yielding 53 nonartifactual ICNs with 490 time points each (for more details, see Datasets in Sec. 4). Therefore, for each subject we have 53 × 53 correlation matrices as features. Because these matrices are symmetric, we extract the nondiagonal upper triangular matrix values (1378 features) for each subject to train the regression models.

### Sampling methods and metrics

As the data need to be distributed across six sites, we employ three strategies to split the data equally. We employ a 90%-10% train-test split at each site. In the first approach, we randomly partition the data into these sites, and within each site, the data is further randomly split into training and testing datasets. This approach is referred to as *random sampling*. In the second approach, we use stratified sampling based on subject age to partition the data into six sites. Within each site, the training and testing datasets were randomly split. We call this *age-stratified sampling*. In the third approach, we group the subjects into different bins based on their age ranges, label these bins and use these labels to perform stratified sampling. We refer to this method as *age-bin-stratified sampling*. All experiments were repeated five times, including splitting the data and training models. To build a centralized model, training data from all the sites is pooled into one training dataset, and testing data from all the sites is combined to create the testing dataset.

We use root mean square error (RMSE) and mean absolute error (MAE) as metrics to measure the performance of the fitted regression models. RMSE is a standard measure used to analyze the performance of regression models. It is a measure of the distance between the predicted values and actual values and is computed as follows:

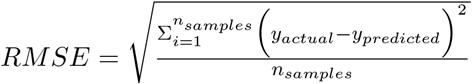

We also compute MAE as it reflects the performance of brain age gaps and is commonly used in literature. It is an average measure of the magnitude of prediction errors without considering their error directions:

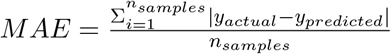

The results show that the decentralized models have similar performance compared to centralized models (see Table 1). For models trained using FreeSurfer features, the decentralized models showed higher RMSE and MAE values on the training data across all sampling strategies, whereas the centralized models demonstrated slightly higher values on the test data. For models trained with GM features, we observed slightly higher error values for decentralized models compared to centralized models on both the training and testing sets across all sampling strategies. For models trained with FNC features, the observations were similar to the ones found on models trained with FreeSurfer features. The decentralized models have slightly higher error values compared to centralized models during training, whereas the testing metrics are slightly improved compared to centralized values.

**Table 1:** Performance comparison of decentralized and centralized models for BAE.

| *Input – data* | *Sampling* | *Method* | *RMSE* |  | *MAE* |  |
| --- | --- | --- | --- | --- | --- | --- |
|  |  |  | train | test | train | test |
| FreeSurfer | random | Decentralized | $3.673 \pm 0.045$ | $3.624 \pm 0.242$ | $3.139 \pm 0.068$ | $3.083 \pm 0.221$ |
| | | Centralized | $3.518 \pm 0.025$ | $3.845 \pm 0.149$ | $2.863 \pm 0.017$ | $3.244 \pm 0.161$ |
| | age stratified | Decentralized | $3.701 \pm 0.052$ | $3.7 \pm 0.184$ | $3.156 \pm 0.051$ | $3.176 \pm 0.172$ |
| | | Centralized | $3.501 \pm 0.012$ | $3.911 \pm 0.137$ | $2.85 \pm 0.01$ | $3.312 \pm 0.159$ |
| | age-bin stratified | Decentralized | $3.652 \pm 0.033$ | $3.646 \pm 0.071$ | $3.131 \pm 0.038$ | $3.102 \pm 0.03$ |
| | | Centralized | $3.522 \pm 0.01$ | $3.862 \pm 0.15$ | $2.863 \pm 0.013$ | $3.24 \pm 0.115$ |
| GM | random | Decentralized | $2.652 \pm 0.132$ | $2.459 \pm 0.156$ | $2.172 \pm 0.134$ | $1.983 \pm 0.147$ |
| | | Centralized | $2.138 \pm 0.015$ | $2.213 \pm 0.141$ | $1.686 \pm 0.012$ | $1.769 \pm 0.109$ |
| | age stratified | Decentralized | $2.588 \pm 0.035$ | $2.489 \pm 0.141$ | $2.063 \pm 0.031$ | $2.029 \pm 0.105$ |
| | | Centralized | $2.151 \pm 0.01$ | $2.185 \pm 0.081$ | $1.691 \pm 0.009$ | $1.761 \pm 0.055$ |
| | age-bin stratified | Decentralized | $2.625 \pm 0.061$ | $2.613 \pm 0.087$ | $2.131 \pm 0.061$ | $2.12 \pm 0.056$ |
| | | Centralized | $2.149 \pm 0.004$ | $2.2 \pm 0.062$ | $1.694 \pm 0.005$ | $1.752 \pm 0.054$ |
| FNC | random | Decentralized | $7.497 \pm 0.076$ | $7.486 \pm 0.074$ | $6.285 \pm 0.078$ | $6.284 \pm 0.066$ |
| | | Centralized | $7.21 \pm 0.005$ | $7.857 \pm 0.084$ | $5.722 \pm 0.009$ | $6.535 \pm 0.06$ |
| | age stratified | Decentralized | $7.502 \pm 0.034$ | $7.494 \pm 0.073$ | $6.298 \pm 0.04$ | $6.284 \pm 0.058$ |
| | | Centralized | $7.203 \pm 0.01$ | $7.968 \pm 0.123$ | $5.709 \pm 0.018$ | $6.623 \pm 0.138$ |
| | age-bin stratified | Decentralized | $7.511 \pm 0.056$ | $7.501 \pm 0.047$ | $6.318 \pm 0.045$ | $6.279 \pm 0.038$ |
| | | Centralized | $7.194 \pm 0.005$ | $7.997 \pm 0.096$ | $5.708 \pm 0.01$ | $6.624 \pm 0.059$ |

Fig. 4 shows the performance of the final (decentralized) model compared to the local models across five runs for different feature sets. Fig. 4(a) and 4(b) show the performance of distributed and local models for FreeSurfer and GM models for 1417 subjects, and Fig. 4(c) shows the performance of the FNC dataset with 11754 subjects. These results show that as the amount of training data increases, the distributed model achieves better accuracy compared to any of the locally trained models.

**Figure 4:**
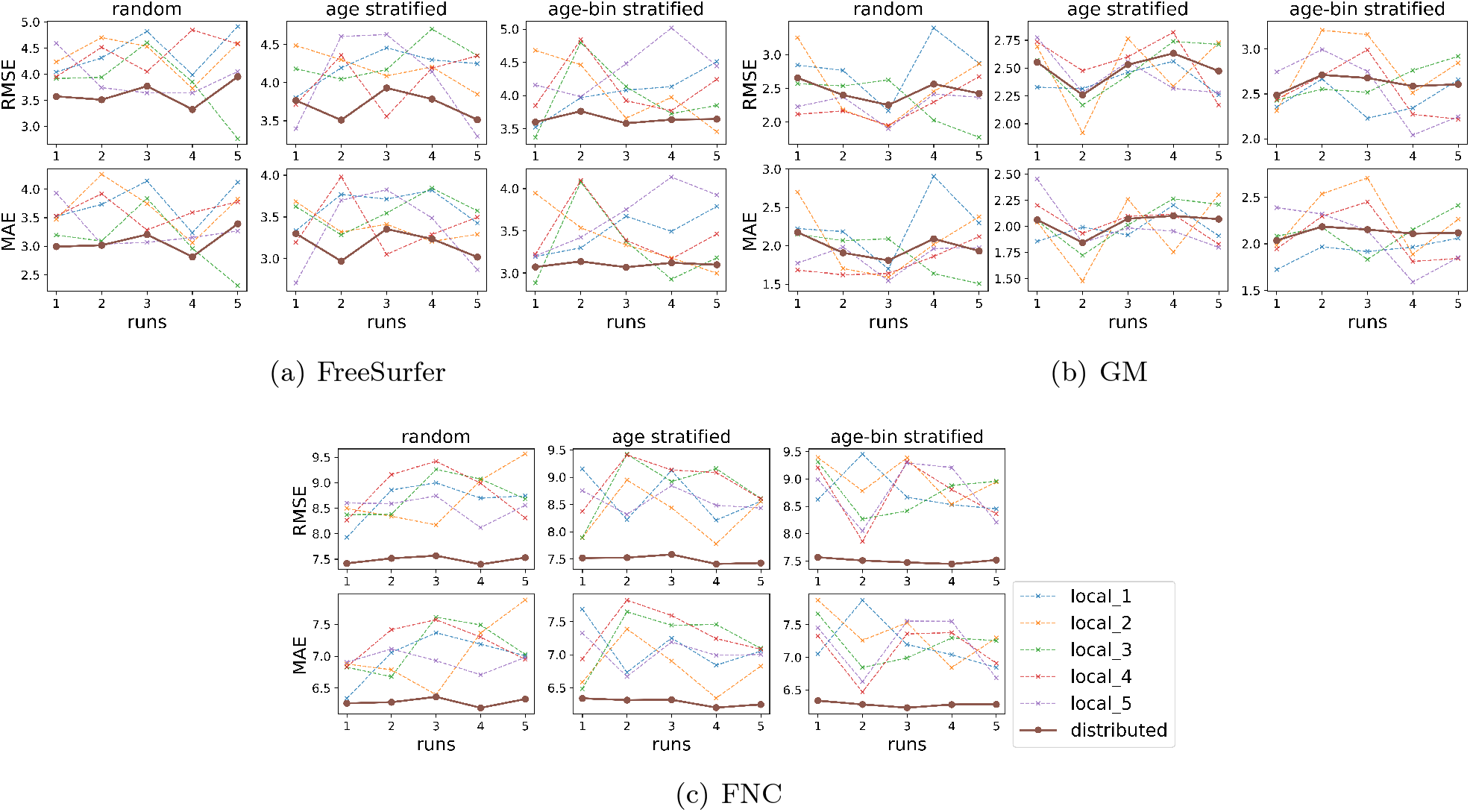
Performance of decentralized models compared to local models across five runs (repetitions) for different feature sets.

We statistically compare the performance of centralized and decentralized models for both measures using a Wilcoxon signed-rank test [34]. This is a nonparametric pairwise comparison test, with no assumptions on the data distribution and a null hypothesis that the differences between two samples have a distribution centered about zero. For each metric (RMSE and MAE), we compare the training scores of decentralized and centralized models for different sampling methods separately using different input data features. We repeat this setup and also compare the testing scores for these metrics. In all the cases, this test fails to reject the null hypothesis indicating that the decentralized models achieved performance similar to that of centralized models. This is a very interesting observation as we tested decentralized models with samples of over one thousand and over 11 thousands, and in all the cases decentralized models perform similar to centralized models. This indicates the de-centralized models work consistently with centralized models for smaller or larger data samples.

## 5 Conclusion

In this work, we employ decentralized machine learning for brain age estimation and compare the results with a centralized model. The decentralized model is trained by utilizing information from locally trained models at different sites and involves no data sharing. Results from models trained on three different feature sets with three different data splitting strategies show that the predictive performance in the decentralized model is consistently better than individual sites separately, and as good as the centralized model. The key benefit of decentralization is that it does not require data sharing and therefore encourages collaboration by allowing different research groups to readily participate in larger studies without worrying about their data-sharing policies or data transmission. Future work will include developing a multishot version of this pipeline coupled with differential privacy strategies.

## Acknowledgments

This work was funded by the National Institutes of Health (R01DA040487), National Institute on Drug Abuse (R01DA049238) and the National Institute of Mental Health (R01MH121246). We sincerely thank Debbrata Kumar Saha and Biozid Bostami for their comments.

## Author Contributions

All the authors helped improve the manuscript. SB designed decentralized models for FNC features, performed the data analysis for all the models and wrote the initial manuscript. RR and HG designed the decentralized model for FreeSurfer and GM features. BR designed feature extraction strategies. JL provided guidance about feature extraction and decentralized model design. EV manages the COINSTAC project and helped write the paper. VDC supervised all the stages of the project and also funded the project.

